# Context-dependent low-dimensional neural dynamics unfold in distinct subspaces, dimensionality, and dynamical strength for natural walking and reaching

**DOI:** 10.64898/2025.12.19.695640

**Authors:** Alissa S Ling, Michael P Silvernagel, Stephen E Clarke, Elizabeth J Jun, Muhammad U Abdulla, Paul Nuyujukian, the Brain Interfacing Laboratory

## Abstract

Awake behaving animal experiments paired with multichannel electrode recordings have advanced motor systems neuroscience in creating models of how the mammalian brain controls movements. However, growing theoretical and experimental evidence question the generalizability of such findings from constrained studies to ambulatory behavior, highlighting a limitation in our understanding of how the brain controls movement. To address this question, spiking neural activity during highly-practiced, routine movement (walking) and goal-directed behavior (reaching towards food) were compared in an unconstrained setting. Kinematic trajectories of the contralateral arm during reaching and walking were statistically similar, as were the average single-neuron firing rates during these respective movements. However, the dimensionality of reaching was higher than that of walking and existed in largely non-overlapping subspaces. Further, when modeled as dynamical systems, reaching decayed 3-5 times more quickly than walking. Taken together, these findings demonstrate that the low-dimensional structure of motor cortex is more complex for goal-directed reaching than in highly-practiced natural movements. Since this difference is primarily observable at the state and dynamical systems level, these findings suggest behavioral context plays a significant role in the coordination of otherwise kinematically similar movements, providing indirect evidence for non-cortical circuits such as central pattern generators.

## Introduction

Understanding how the brain controls diverse movements in natural environments would enable the development of novel therapeutics to restore behavior and function after injury. Decades of research in motor systems neuroscience involved awake, behaving animal experiments with multichannel electrode arrays to elucidate the neural mechanisms behind voluntary movement in the mammalian brain. Rhesus macaques have been a common model species because of their neuro-anatomical sim-ilarities to humans McCowan et al (2008); Bizzi et al (2000) and their ability to perform a wide range of simultaneous naturalistic behaviors, such as arm reaching Churchland et al (2012), walking Foster et al (2014), foraging Eisenreich et al (2019), and social behavior Testard et al (2024). A common behavioral paradigm to investigate how the brain controls goal-directed movement is a “center-out” reach task where an animal reaches towards targets on a screen placed in front of them Georgopoulos et al (1982). Neural data is captured from a direct, wired connection to the implanted array and kinematic data of the reaching arm is captured by tracking the hand using an infrared bead. The animal is constrained except for the movement of interest which helps isolate the desired behavior and reduce confounding variables such as tail or leg movement to draw tighter correlations with the brain activity. These traditional experiments are foundational for understanding how the brain controls reaching behavior at the single neuron and population level.

Classical experiments with chair-seated monkeys have shown that motor cortex is correlated with dynamic variables of movement, such as action value, vigor, and speed by comparing firing rates and tuning curves of neurons Grillner (2002). By contrast, other studies suggested that individual M1 neurons controlled the direction of reaching according to high-level movement goals rather than through direct muscle activation Georgopoulos et al (1986). Recent studies applying control theory and multidimensional statistical estimation tools to firing rates have uncovered relationships between high-dimensional population activity of neurons and behavior that cannot be seen on a single-neuron level or trial-averaged basis Churchland et al (2012). These findings have revealed rich, low-dimensional neural dynamics that lie within a preserved neural manifold underlying multiple reaching behaviors Gallego et al (2018); Perich et al (2025); Pandarinath et al (2015). Consequently, one goal of investigating low-dimensional neural structure could be to describe the complete neural manifold of cortical activity for a full repertoire of behaviors.

Accomplishing this goal requires studying more complex and unconstrained behaviors, as simpler, constrained tasks may be artificially limiting neuronal task complexity Shadmehr and Wise (2004); Gao and Ganguli (2015). Theoretical evidence suggests that the variability of recorded neuronal activity may also increase with task difficulty, possibly changing the dimensionality of neural manifolds constructed from constrained studies Gao and Ganguli (2015); Sompolinsky (2014). Thus, a key unanswered question is how consistent and generalizable observed neural dynamics are between constrained and freely-moving, natural behaviors. In recent years, freely-moving animal studies have begun to answer this question, collecting a rich repertoire of behaviors and contexts Mimica et al (2023); Foster et al (2014); Capogrosso et al (2016); Berger et al (2020). Evidence in rodent studies comparing similar behaviors performed under different environments (i.e., constrained versus unconstrained) showed that motor cortex plays a more complex role than exclusively controlling muscle movements Miri et al (2017). This study showed that optogenetically increasing inhibition in caudal forelimb area disrupted the execution of a precision pull task but not a treadmill task. This may suggest that motor regions have different levels of involvement for walking and reaching. Prior unconstrained macaque studies explored walking on a treadmill Foster et al (2014, 2011); Presacco et al (2011); Fitzsimmons et al (2009) or through a narrow corridor Capogrosso et al (2016). Findings from these freely moving platforms suggest that different behaviors, such as engaging multiple limbs versus one limb, give rise to different neural activity Fitzsimmons et al (2009); Mori et al (2006). In freely-moving social behavior, the structure of neural population activity was context dependent for both the movement of the animal and the social context Testard et al (2024). Taken together, these studies suggest that the context of a behavior changes the involvement and neural population structure in motor cortex.

One approach to further investigating the generalizability of neural dynamics in different behavioral contexts is through performing different behaviors in an unconstrained freely-moving environment. This study aims to address the role of M1 in unconstrained reaching and walking in primates and whether similar movements performed under different contexts change cortical activity. Two similar kinematics behaviors (reaching and walking) in slightly different contexts (goal-directed vs. routine) were compared to characterize the low-dimensional neural dynamics of unconstrained structure. The study provides a behavioral paradigm in a more natural environment than treadmill studies, but still captures task-based and repetitive, stereotyped movements Foster et al (2014). This investigation is also a potential way to explore central pattern generators in large animals more closely related to humans than other animals with rhythm-generating circuits, such as cats Kiehn (2016); Drew (1993); Sherrington (1893).

This study collected free behavior in macaques in an experimental framework that captures wireless neural data and its relationship to full-body movement Silvernagel et al (2021). This freely-moving environment captured behaviors of greater complexity than those seen in a constrained setting due to the coordination of multiple limbs, increased visual and sensory information, and different goals. From this repertoire, we compared the neural response of two similarly kinematics unconstrained arm movements in different contexts: arm swing while walking and arm reaching to food. The key difference between these task conditions is the movement intention, where unconstrained reaching is an effortful, directed movement towards a salient reward, whereas unconstrained walking is an highly-practiced, repetitive, and stereotyped cycle with the intent of moving forward. We sought to answer how the context behind similar behaviors influences neural activity in M1 on both a per neuron basis and also at the neural population level for the gait cycle and reaches to food. This study provides insight on how the primate brain may control free ambulation under different behavioral contexts.

## Results

Two adult male rhesus macaques, Monkey U and Monkey C, performed hundreds of trials over multiple days of natural, free walking and reaching, time-synchronized with neural data in a large observational enclosure (Figure 1a) Silvernagel et al (2021). Figure 1b displays example arm kinematics of the shoulder, elbow, and wrist for the swing, stance, and subsequent arm reach for Monkey C. The example kinematics represent the behavioral task, as the monkey completes a gait cycle and then reaches for food in a bowl in one smooth movement. Analysis of forelimb movement kinematics were performed to compare the kinematics of the walking and reaching contexts. Figure S4 displays the 2D center of mass kinematics of each monkey for three experimental days as they perform the behavioral walk-to-reach task. These three days were representative of the kinematics for the other behavioral days.

**Fig. 1.**
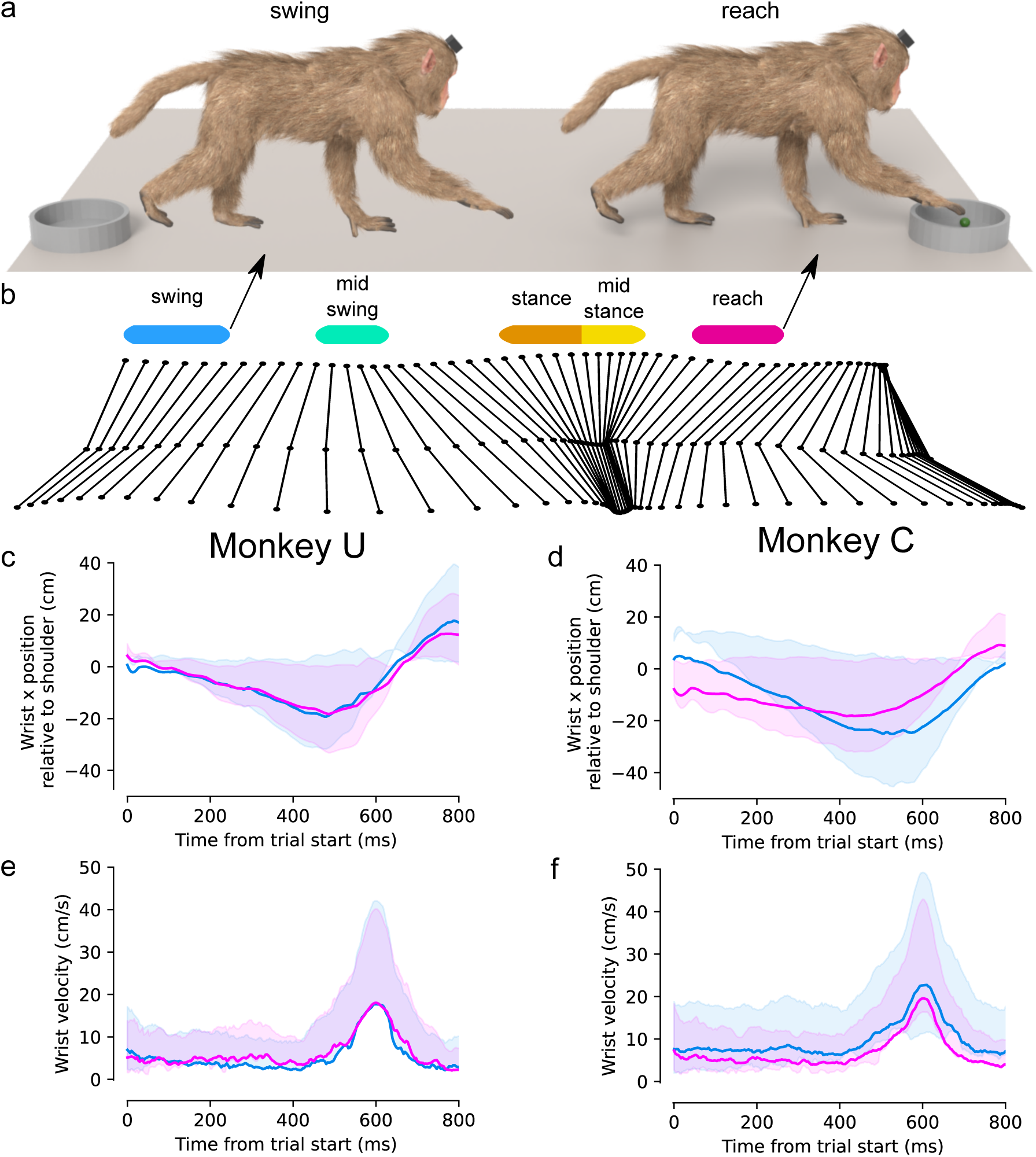
Behavioral task and kinematic comparison of walking versus reaching. a) Diagram of the walk-reach behavioral task, where animals take 1-3 gait phases per lap before reaching towards food in a bowl. After a reach is complete, the monkey turns around and walks to the bowl on the opposite side of the observational enclosure **?**. b) Continuous arm kinematics of the shoulder, elbow, and hand for Monkey C for one example lap. Kinematics were extracted from 4 RGB-D cameras, sampled at 30 Hz using an automated pose-tracking algorithm **?**. Each of the colored bars represent 200 ms of time starting at the beginning of each gait phase: swing (blue), midswing (teal), stance (orange), midstance (yellow), and reach (magenta). The blue bar spans 10.7 cm. The arrow from the blue bar points to the monkey in the swing phase, and the arrow from the magenta bar points to the same monkey in the reach phase to highlight the two behaviors that have similar arm kinematics, under different contexts. c) The position of left wrist kinematics relative to the neck in the forward dimension for the 800 ms of the reach towards food and swing phase for Monkey U is plotted. The three-dimensional position was manually hand-tagged for all point cloud frames at the neck and the wrist. Solid lines represent the median for all hand-tagged trials, while shading represents the 25% and 75% percentile ranges. d) Same as in c), but for the right wrist of Monkey C. e) The velocity of the left wrist relative to the shoulder is plotted for each trial for Monkey U. Each trial is aligned to the peak velocity. Stance phase occurs when the wrist is at the lowest velocity, between times 0 and 200 ms, and midstance occurs at 200 to 400 ms. Swing and reach onset occur at 400 ms, gain peak velocity at 600 ms indicating the start of midstance, and end at 800 ms either towards a new stance trial or grasp to food in a bowl. f) Same as in e), but for the right wrist of Monkey C.

Figures 1c and d quantify the contralateral wrist position in one dimension relative to the neck. Solid lines represent the average swing (blue) and reach (magenta) across all trials for three hand-tagged days for Monkey U and C, respectively. The 25% and 75% percentiles of the swing and reach are shown as lightly shaded regions around the median. The median contralateral wrist velocity for the walk (blue) and reach (magenta) were derived from the hand-tagged wrist positions. Figures 1e and f plot the statistically similar velocity profiles for both swing and reach for both monkeys. Table S2 and Table S3 further quantify kinematic variables such as maximum velocity, maximum acceleration, minimum acceleration, millisecond trial start time, trial end time, and peak velocity time. For Monkey U, all six kinematic features were statistically indistinguishable with p *<* 0.05 after performing a Mood’s median test for the swing compared to the reach. For Monkey C, all features but the maximum velocity of swing versus reach were statistically indistinguishable with p *<* 0.05 using a Mood’s median test.

Neural data were recorded from a 96-channel electrode array (Blackrock Microsystems, Salt Lake City, UT) implanted in M1 right arm cortical region for Monkey U and M1 left arm cortical region for Monkey C. Neural data were wirelessly telemetered with a battery-powered transmitter (Cereplex W, Blackrock Microsystems, Salt Lake City, UT). Figures 2a and b show the median firing rates sorted by the most active electrode for all swing and reach trials. Figures 2c and d show 1-1.5 seconds of binned neural threshold crossing rasters for Monkeys U and C for the contralateral limb. Both rasters show modulation during 200 ms periods of swing (blue) and reach (magenta) for the contralateral arm. A two-sided Kolmogorov-Smirnov test was performed to assess whether the observed swing and reach median firing rates were drawn from the same distribution. For both monkeys, neuron firing rates were not significantly higher in the reach condition than the swing condition (Monkey U p = 0.96, Monkey C p = 0.65). This suggests that motor cortex has similar levels of activity during directed reaches and while walking.

**Fig. 2.**
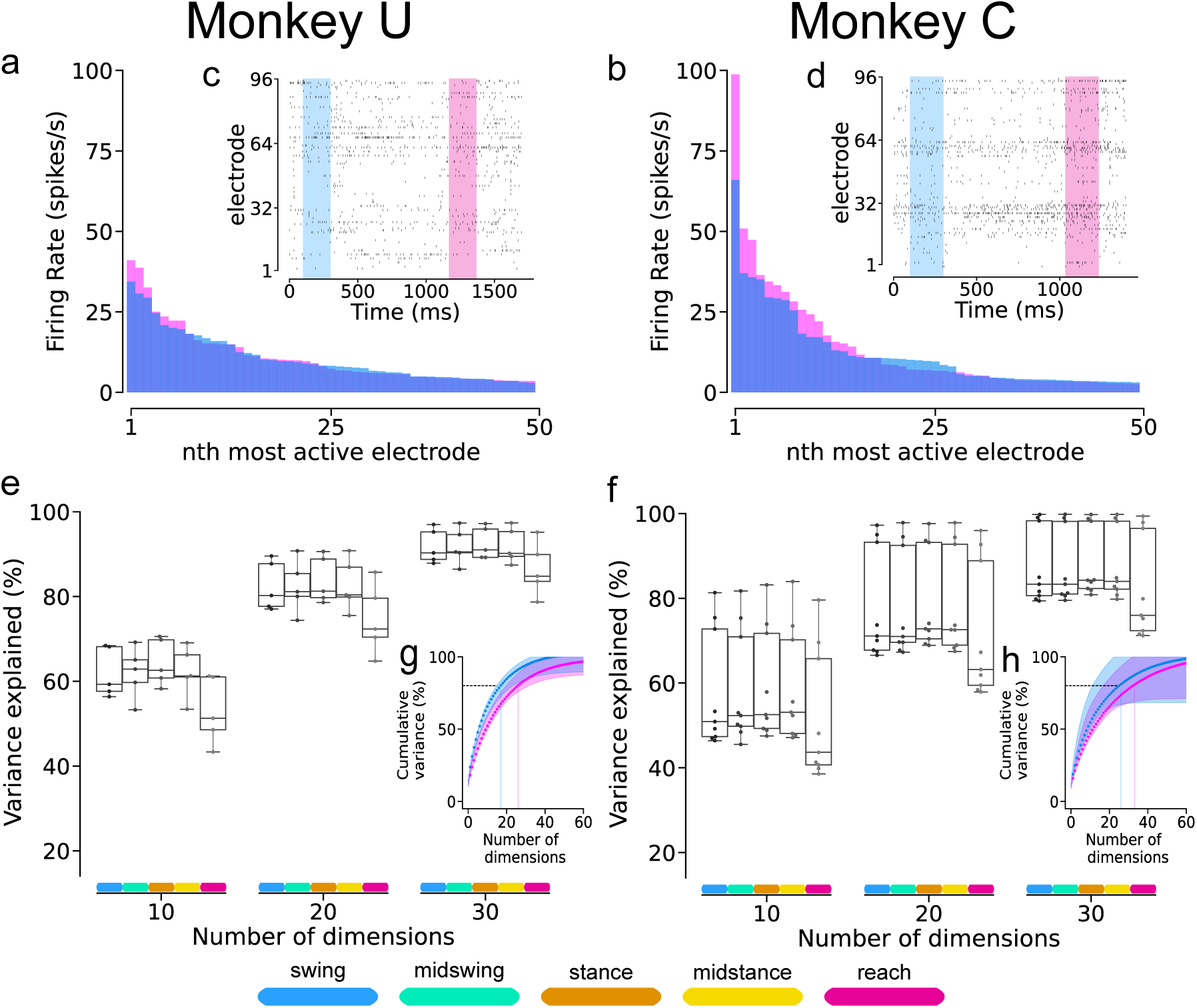
Wireless neuron population recordings from primary motor cortex during ambulatory walking versus reaching are similar in firing rates, dimensionality, and power distribution. a) Trial averaged firing rates from left M1 arm area comparing swing (blue) and reach (magenta) conditions for Monkey U. Electrodes were sorted from highest to lowest firing rate for each condition, and 50 out of 96 electrodes are displayed, as some have negligible activity. b) Same as in a), but for Monkey C in right M1 arm area. c) Neuron spike raster plots for Monkey U from M1 array in left arm regions spanning 2 seconds of data from 96 electrodes. Vertical bands span 200 ms and are color coded for an example swing (blue) and reach (magenta) trial. d) Same as in c) but for Monkey C (right M1 array) spanning 1.5 seconds, as Monkey C performed the walk-to-reach task faster than Monkey U. e) Individual fractions of variance for the top 10, 20, and 30 PCs accounted for the swing, midswing, stance, midstance, and reach epochs in it owns subspace for each day for Monkey U. Each point is the fraction of variance for each day and the box and whisker plots show the distribution over multiple days. f) Same as in e) but for Monkey C. For both monkeys, all four gait conditions contribute similar fractions of variance in its own subspace, suggesting that phases of the gait cycle have similar dimensionality. The reach epoch contributed significantly less fractions of variance, showing that the reach is higher dimensional than the gait cycle, and that different contexts change the dimensionality in M1. g) Median and inter quartile ranges across experimental days for the cumulative number of PC dimensions needed to capture a fixed percentage of variance for the swing (blue) and reach (magenta) conditions. The dotted vertical line indicate the number of dimensions needed for 80% of the variance. For Monkey U, 17 dimensions for the swing phase and 26 dimensions for the reach phase explained 80% of the variance. Both a two-sided and greater Kolmogorov-Smirnov test were performed on the median cumulative variance to test if the distributions of the swing and reach eigenspectra come from the same distribution and if the swing eigenspectra is statistically greater than the reach. For Monkey U, p *<* 0.001 for a two-sided KS test showing that they are statistically different distributions, and p *<* 0.001 for a greater KS test showing that the reach eigenspectra is statistically smaller than the swing eigenspectra. h) Same as in g), but for Monkey C with 24 dimensions for swing and 33 dimensions for reach. For Monkey C, p = 0.24 for a two-sided KS test and p = 0.12 for a greater KS test.

The dimensionality of neural activity for swing and reach were compared to investigate the variance associated with both behaviors (reaching vs walking). Similar spectral distributions of variance would indicate a similar dimensionality of reaching and walking. On the other hand, differing spectral distributions would suggest that the behavioral context altered the dimensionality in M1. Principal component analysis was used to create two subspaces for the same number of swing and reach trials for each experimental day. Figures 2g and h show that the median cumulative eigenspectra are separable between the two behaviors for both monkeys. Further, a greater number of dimensions are needed to account for the same fraction of variance in reaching than in walking. For Monkey U, 17 dimensions for the swing phase of walking and 26 dimensions for reaching are needed to explain 80% of the total variance. For Monkey C, the number of dimensions needed are 24 and 33, respectively. Consistent with the separable PCA eigenspectra for swing and reach conditions, linear discriminant analysis shows that they are well separated, indicating that the behaviors can be readily distinguished in low-dimensional neural spaces (Fig. S5).

Further analysis was performed to compare the variance of the full gait cycle to the reach (Figures 2e and f). The gait cycle was separated into four phases, swing (blue), midswing (green), stance (orange), and midstance (yellow), consisting of 200 ms per phase time aligned to the velocity profiles from Figure 1e and f. As above, the reach trials remained the same, and the number of walking trials matched the number of reach trials per experimental day. Individual fractions of variance were calculated by projecting the neural activity for each phase into its own subspace for the top 10, 20, and 30 PCs. When projected into its own subspace, the four phases of gait explained similar fractions of variance compared to each other, while the reach explained significantly lower percentages of variance than any of the gait phases. This result is consistent with the finding that reaching is higher dimensional than walking.

The fractions of variance shared between walking and reaching were investigated by cross-projecting each epoch into another epoch’s subspace in the top 10 PCs (Figure S6). The reach neural data contributed less than 21% of variance to every gait subspace, while every gait neural epoch contributed less than 25% variance to the reach subspace for both Monkey U and C. This finding demonstrates that the reach and gait subspaces have very little overlap in state space. When comparing the contributions of variance between the swing and stance phases of the gait cycle, the swing and midswing subspaces captured less than 20% of the variance of the stance epoch. Similarly, the stance subspace captured less than 26% of the variance of the swing epoch. These results suggest that both contextually different yet kinematically similar behaviors and kinematically different behaviors (swing vs. stance or reach vs. stance) lie in separate subspaces. Taken together, these findings suggest that spiking neural spectral power and extent of coordination between neurons are different between walking and reaching.

Low-dimensional neural dynamics of walking and reaching were investigated next to describe how M1 activity evolves over time to control these unconstrained behaviors Churchland et al (2012); Yu et al (2007). A low-dimensional subspace was constructed from all gait cycle trials (200 ms of swing, midswing, stance, and midstance each) for each day using PCA. The number of gait trials per day is displayed in Table S1. Both the gait trials and reach trials were then projected onto this resultant subspace. Trial-averaged neural trajectories in state space for the four phases of the gait cycle (swing, midswing, stance, and midstance) and reaching are displayed in Figure 3a and b for one example experimental day (U221216 01 and C220422 01). The two panels show the same neural trajectories from a different view and show the separation of the reach from the gait cycle. For both monkeys, the gait cycle has cyclic trajectories, while the reach was separate and shifted in state space from the oscillatory direction of walking. Figure 3c and d plot the distribution of the median distance in the top 10 dimensions between every point in a 200 ms reach trial to every point in a 200 ms swing, midswing, stance, and midstance trial for all trials across all days. The results show that the reach trajectory is spatially closest to that of the swing or midswing phase (significance of p *<* 0.001 using a ranksum test), the most kinematically similar behavior to reach in state space. This finding was consistent for both monkeys for 10, 40, and 96 dimensions (Figure S7).

**Fig. 3.**
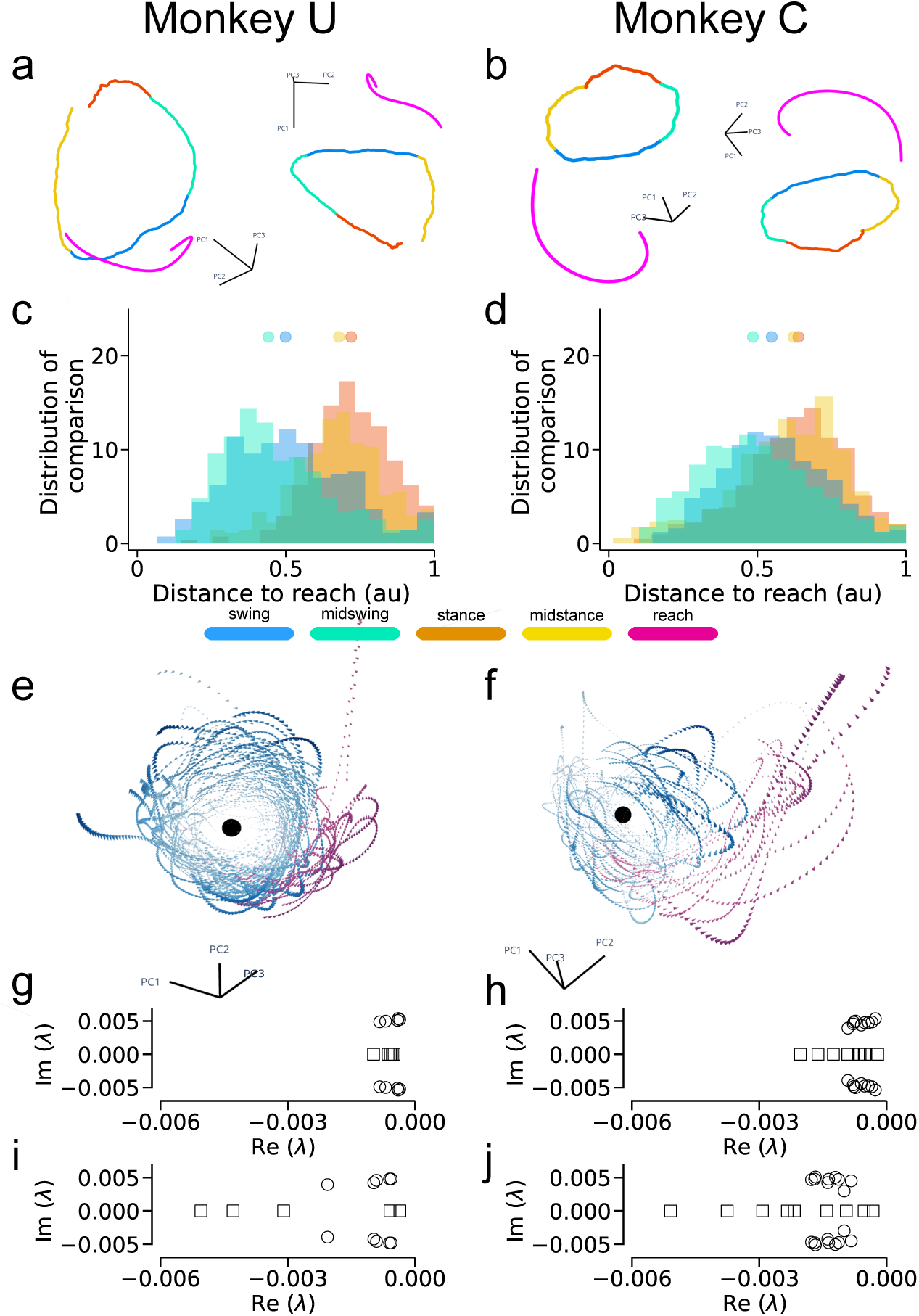
Low-dimensional neuronal dynamics of walking versus reaching evolve into separable and shifted dynamics. a) Neural trajectories of walking and reaching in 3-dimensional neural state space constructed using PCA for Monkey U. The gait cycle has oscillatory dynamics, and the reaching trajectories are closest to that of the swing phase in PC space. The average trajectory for all trials for each phase (swing 300 ms, midswing 200 ms, stance 300 ms, midstance 400 ms, reach 600 ms) is plotted and color coded as defined by the colorbars. Epoch times are chosen for for visualization purposes. b) Same as in a), but for Monkey C. c) The histogram calculates the median closest distance in 10-dimensional PC space for each gait phase to each reach trial for 200 ms for Monkey U. Each dimension is scaled by the fractional variance contributed, and the distance values were scaled to be between 0 and 1 using maximum absolute scaling. d) Same as in c) but for Monkey C. For both monkeys, swing and stance median distances to the reach were statistically different using a Mood’s median test with p*<*0.001. e) Neural dynamics of the gait cycle exhibit oscillatory, counterclockwise behavior converging towards a fixed point in the center. All gait cycle trials (blue) and all reach trials (magenta) projected into the gait dynamical system are plotted for one experimental day for Monkey U (U221130 01). Cones show the direction of oscillation and the black circle represents the fixed point of the system. f) Same as in e) but for Monkey C (day C220406 01). g) The top three eigenvalues of the dynamic A matrix for the gait cycle are plotted for Monkey U. The first real eigenvalue (squares) quantifies the decay or growth factor, and first pair of imaginary eigenvalues (circles) quantify the rotational component. The top three dynamical eigenvalues always have a negative real component showing that the neural dynamics of the gait cycle are stable and decaying towards a fixed point in the center of the cycle. h) Same as in g) but for Monkey C. i) The top three eigenvalues of the dynamic A matrix for the reach are plotted for Monkey U. The eigenvalues for the reach dynamic matrix also has a negative, real eigenvalue (square) and a pair of negative imaginary eigenvalues (circles).The ratio of the median of the real reach eigenvalues to the median of the real gait eigenvalues is 5.3, and the ratio for the imaginary eigenvalues is 2.3. The real eigenvalues for the reach dynamics are much larger than the real eigenvalues for the gait dynamics suggesting that the neural reach dynamics converge to a fixed point more rapidly than the gait dynamics. j) Same as in i) but for Monkey C. The ratio of the median of the real reach eigenvalues to the median of the real gait eigenvalues is 3.3 and 2.2 for the imaginary eigenvalues.

The low-dimensional neural dynamics of walking and reaching were modeled as two separate linear dynamical systems to quantify the oscillatory nature of the two systems. PCA trajectories of each behavior were fit to unique linear dynamical systems using least-squares to approximate the change in low-dimensional neural trajectories over time. Figures 3g and h plot the first real eigenvalue (squares) and the first pair of imaginary eigenvalues (circles) for walking for each day for Monkey U and Monkey C. Likewise, Figures 3i and j plot the same, but for reaching. The real eigenvalues of *A_gait_* and *A_reach_*were always negative, indicating that these dynamic systems for walking and reaching are stable and decaying towards a fixed point. Comparable, non-zero complex eigenvalues of *A_gait_* and *A_reach_* show that both systems exhibit similar degrees of oscillatory dynamics. An example of the neural gait and reach trajectories fit to the gait dynamical system is shown in Figures 3e and f for one experimental day per monkey (U221130 01 and C220406 01). The magnitude of the real eigenvalues of reaching were proportionally much larger than that of walking. For Monkey U, the ratio of the median of the real reach eigenvalues to the median of the real walking eigenvalues was 5.3× larger, while for the Monkey C, it was 3.3× larger. This suggests that neural reach dynamics decay more rapidly towards a fixed point than the dynamics for walking. In summary, reaching is higher dimensional and has faster temporal dynamics, while and walking is lower dimensional and slower.

## Discussion

This study presents a dynamical systems perspective of natural walking and reaching behavior in freely-moving rhesus macaques to evaluate the impact of behavioral context (directed versus routine) on motor cortical neural activity. The results support that the behavioral context changes the correlational structure between neurons, the low-dimensional neural manifold, and the dynamical systems complexity. The position, velocity, and acceleration of reaching and walking were statistically similar, enabling the comparison of similar movements performed under different contexts. The single-neuron responses in motor cortex were not statistically different, as shown by the similar median distributions of firing activity across channels and trials. Specifically, Monkey U had similar kinematics of reach and swing, which resulted in a similar distribution of neural firing rates across all trials. Of note, Monkey C’s peak swing velocity was higher than his peak reach velocity, which may have reflected in slightly, but not significantly, higher firing rates during reaching. This may show a relationship between peak velocity and firing rates, which could be examined in future work that explores how varying speed and movement direction correlates with neural activity.

Differences in neural activity between similar kinematics were primarily apparent in low-dimensional state and and dynamics level. The neural dimensionality of walking was lower than reaching suggesting that M1 is less involved at a population level during highly-practiced walking. Low-dimensional neural population activity in M1 operated in unique and non-overlapping subspaces where reaching was shifted from walking in state space. When modeled as linear dynamical systems, the dynamical eigenspectra of walking and reaching both had negative imaginary eigenvalues exhibiting converging, oscillatory dynamics. However, the top real eigenvalue for reaching was 3-5 times larger than that for walking, demonstrating that reaching exhibited a stronger decay factor towards its fixed point. The dynamical eigenspectra for walking had less-damped, slower-decaying oscillatory dynamics. It may be strategically useful for motor cortex to have a stronger drive to collapse during transient reaching movements, as a system with faster decaying temporal dynamics can stop more quickly. Likewise, it may also be useful for walking to be less damped, because walking is more likely to be continuous for longer periods of time. These results potentially provide correlative evidence for a central pattern generator in primates, as they suggest that motor cortex is less involved in walking and more involved in driving goal-directed behavior such as reaching.

This work builds on previous, partially constrained animal studies Foster et al (2011, 2014); Gilja et al (2010); Borton et al (2013); Schwarz et al (2014); Berger et al (2020); Capogrosso et al (2016); Pre-sacco et al (2011); Fitzsimmons et al (2009) and further demonstrates a freely-moving rhesus model with which to explore generalized low-dimensional motor neural dynamics. In previous treadmill studies, animals were limited to a single movement axis and walking speed was externally controlled by a treadmill Foster et al (2014); Berger et al (2020). In this study, walking speed was volitional and varied by the monkey, and had more dimensions of freedom to move around in the observational enclosure, as seen in the variability of the center of mass trajectories through the space. This study confirms that neural dynamics during walking are oscillatory, a characteristic found in a previous treadmill study Foster et al (2014). Future work may further characterize the relationship between state space trajectories and speed in a naturalistic environment to develop brain machine interfaces for natural walking.

Relating back to classical, constrained reaching tasks, our results provide evidence that some insights about motor cortical control from constrained environments may generalize to unconstrained environ-ments, such as the rotational structure of low-dimensional neural activity during reaching Churchland et al (2012). However, our results show that different behavioral contexts have separate eigenspectra, subspaces, and dynamics of cortical activity; suggesting that additional work is needed to identify a single, generalizable model for motor cortical structure. This supports the hypothesis that sim-pler, constrained movements are not capturing the full repertoire of neural activity and may be artificially constraining observed neural manifolds Gao and Ganguli (2015). Indeed, recent work exploring different behavioral contexts and motor tasks have also found that M1 is higher dimen-sional and more flexible than previously observed Amematsro et al (2025). Exploring a wide range of different unconstrained primate behaviors may provide a more comprehensive understanding of the high-dimensional motor neural population manifold. Future work may seek to directly compare constrained and unconstrained reaching tasks to further elucidate this finding.

This study describes how neural dynamics evolve across different behavioral contexts and provides evidence that motor cortex exhibits varying complexity to control these behaviors. These findings are in line with prior work showing that increasing inhibition in motor regions of mice results in a behavioral deficit for reaching but not for walking Miri et al (2017); Drew (1993). Here we demonstrate similar conclusions about the role of M1 in different contexts in primates. Additionally, previous lesions studies with unconstrained behavior were performed in rodents or cats. Clarification about this context dependence in healthy or injured cortex may involve lesions in freely-moving environments in primates Bray et al (2023). Future lesion studies may help uncover causally necessary brain regions for naturalistic movements, which could lead to novel therapeutics that restore natural function Nudo et al (2003).

## Methods

### Animal Procedures & Behavioral Task

All animal procedures and protocols were approved by the Stanford University Institutional Animal Care and Use Committee (IACUC). Two adult, male rhesus macaques (*Macaca mulatta*; Monkey C and Monkey U) separately performed an unconstrained walk and reach task. Monkey C, age 14, weighing 15.3 kg, was implanted with two 96-channel multielectrode arrays (Blackrock Microsystems, Salt Lake City, UT) in arm regions of primary motor (M1) and dorsal premotor (PMd) cortex determined by visual anatomical landmarks using standard neurosurgical techniques on March 25, 2021. Monkey U, age 12, weighing 14.5 kg, was implanted with three 96-electrode multielectrode arrays (Blackrock Microsystems, Salt Lake City, UT) in arm regions of primary motor (M1) and dorsal premotor (PMd) cortex determined by visual anatomical landmarks using standard neurosurgical techniques on August 4, 2017. Only M1 data was used in this analysis for both monkeys.

An unconstrained reach-walk behavioral task was captured for two macaques in a large observational enclosure similar to those previously developed Silvernagel et al (2021); Bala et al (2020). Task behavior consisted of laps, in which the animals walked between bowls placed on opposite sides of the enclosure, and then stopped and reached for treats. Laps contained different numbers of walking steps, but the average trial had two to three arm swings before the reach at the end of each lap. The reach trials were counted as valid only if the macaque maintained a standing posture on all fours. Experimental sessions and trial counts for Monkey C and Monkey U are summarized in Table 1. Animals voluntarily walked and reached for treats and were free to sit and wander around the observational space.

The gait cycle was defined by four major phases, swing, midswing, stance, and midstance for the contralateral arm. The beginning of stance periods and reach periods were hand tagged from RGB videos for the entire experimental session for each day. Stance periods were defined as the beginning of hand contact with the floor, swing periods were the beginning of hand lift off. Reach periods were when the animal initiated a reach for food in the bowl with the contralateral arm from the implant in M1. Swing periods were inferred from the data, and calculated as the time exactly in the middle bounded between two hand-tagged stance phases. The timing of midstance and midswing were also inferred; midstance was bounded by stance and swing, and midswing was bounded by swing and stance.

### Kinematic feature extraction

Wrist positions were obtained through manually hand-tagging locations of the neck and wrist on individual point cloud frames for all trials for three experimental days per monkey. Hand-tagging was completed in a custom application written in Python that loaded each point cloud for exper-imental sessions U221130 01, U221201 01, U221202 01, C220406 01, C220411 01, and C220422 01.

For Monkey U, a total of 565 walking trials and 238 reaching trials were manually hand-tagged, and for Monkey C, a total of 344 walking trials and 135 reaching trials were manually hand-tagged. The epoch lengths and timing with respect to trial start were obtained from the wrist velocities (Figures 1e and f) with stance between 0-200 ms, midstance from 200-400 ms, swing from 400-600 ms, midswing from 600-800 ms, and reach from 400-600 ms. Swing trials were defined as a 200 ms period beginning at the initiation of wrist lift off during walking, while reach trials were defined as a 200 ms period starting at the initiation of wrist lift off before a reach to food. Quantitatively, the beginning of both the reach and swing were 200 ms before peak velocity (Table S2 and Table S3).

The center of mass of the monkey was extracted from all point cloud frames for each experimental session using the Open3D Python package Zhou et al (2018). Using the RANSAC (Random sample consensus) segmentation algorithm, the floor and background were removed, isolating the monkey’s point cloud. The center of mass was found by calculating the centroid of that point cloud. The center of mass during the walking and reaching behavior was stereotyped with linear kinematic paths between the two food bowls (Figures S4c and d). Trials that were outside of the standard deviation distance from the path were discarded, as this was when the monkey was distracted from the task and freely exploring the observational enclosure.

### Neural data processing

Neural data were recorded wirelessly in arm regions of primary motor cortex using a wireless head-stage (details provided below). These signals were acquired at 30 kHz. Due to the unconstrained wireless transmission of recorded neuronal activity, there was more motion artifact, static, and drops present in the data compared to standard constrained studies. Therefore, standard high pass filtering and thresholding to detect action potentials (spikes) were prone to false positives. We filtered the true spikes from the artifact by using cross correlating potential spikes with a wavelet template taken from our constrained rig and also filtered based on the biological size of the depolarization.

Our wireless neural data contained artifact due to wireless electrical noise from the surrounding observational enclosure and unconstrained head movements. These artifacts are not present in traditional chair experiments, as the animal is head-fixed with a tethered connection to neural data. This artifact was removed by taking a rolling root mean square (RMS) of 100 sample intervals for the entire neural high pass filtered data for that day. Drops in the data had an RMS of less than 5 *µ*V and very large changes due to static causing voltages to sharply max out were and RMS above 70 *µ*V. Neural data was counted as valid if the RMS was between 5 and 70 *µ*V, otherwise, it was not included in the subsequent thresholding. We then took the RMS of 60 seconds from the middle of an experimental day of valid data for each electrode and multiplied by a factor of-4 to obtain a per electrode threshold Chestek et al (2009). Thresholds were around-40 to-50 *µ*V.

Smaller artifacts passed threshold detection, so an additional spike filtering algorithm was used to sort these smaller artifacts from real spikes. We performed a cross correlation between any detected spike and wavelet template taken from the same array in a constrained rig. We counted a potential spike as true if the cross correlation peak was around 47-48 samples, ensuring that the depolarization was in the correct spatial location. Additionally, physiologically the depolarization of a spike should last less than 200 ms which is about 10 samples Scott and Kalaska (2021). We found the minimum of the spike and found the two inflection points around the minimum. Spikes were included if the depolarization lasted between 7 and 14 samples. After the additional spike filtering, a 30kHz raster was constructed from the filtered neural data and then downsampled to a 1kHz raster. Finally, to remove any remaining array-wide artifact, spikes that appeared on more than 30 electrodes (1/3 of the array) within the same millisecond were removed. An example of the spiking activity for each electrode and each trial is shown for Monkey C in Figure S11 and Monkey U in Figure S9. Average peri-stimulus time histograms for the same example day for swing, midswing, stance, midstance, and reach are shown in Figure S12 and Figure S10 for Monkey C and Monkey U respectively. Median firing rates distributions for swing and reach were compared with a rank sum test (Figure 2a and b).

Neural activity was present on all intact channels, whereas channels with no detected activity were either damaged or interfaced with neurons that did not spike. Electrodes with firing rates less than 5 percent of the maximum were excluded before comparing firing rate distributions, resulting in a cutoff of 3.0 Hz which included 25 electrodes for Monkey C, and a cutoff of 1.8 Hz which included 28 electrodes for Monkey U. Although some electrode arrays had a firing rate of 0, state-space and dynamical analyses capture correlational structure where it exists, so including potentially empty rows does not alter the low-dimensional findings Cunningham and Yu (2014).

We then quantified the percentage of dropped data and artifact present for all trials to determine if any correlation existed with a specific gait phase or the reach. Artifact and dropped data did not correlate with any specific trial type or period of time during a trial. Trials with 20% of dropped data for Monkey C and 40% of dropped data for Monkey U were excluded from analysis. All analysis were performed on remaining trials. Experimental sessions, number of swing trials, and number of reach trials are reported in Table S1.

### Synchronization of neural and behavioral data

Neural data were transmitted from the animal through a wireless headstage (CerePlex W, Blackrock Microsystems, Salt Lake City, UT). Outside the enclosure, 16 panel antennas (PA-333810-NF 3.3GHz-3.8GHz 10dBi Panel Directional Outdoor Antenna, FT-RF, Jhubei City, Taiwan) are evenly spaced to receive signals from two wireless headstages operating at distinct frequencies in the 3-4 GHz range. This allows hundreds of electrodes of neural data to be recorded simultaneously with freely-moving behavior. Raw neural data were sampled at 30 kHz and saved to a SQLite database.

The neural data were collected from a neural acquisition system and the behavioral data was collected from four RGB-D Microsoft Azure Kinects at 30Hz Silvernagel et al (2021). The four cameras were synced to each other using a lead camera that controlled the acquisition of other cameras. The cameras were asynchronously triggered by the neural acquisition system, allowing for high fidelity temporal alignment between the distinct data streams. Three square LED lights were placed outside of the observational enclosure on the three sides that did not include the tunnel entrance. The status of the light (on or off) appeared in the RGB frames. The binary status of the light was used to encode the frame number. The sequence of lights created a continuous binary stream with an unique 8-bit header to signify a starting frame.

### Low-dimensional linear analysis

Linear dimensionality reduction techniques map neural firing rates of approximately 10^2^ neurons to an n-dimensional neural state space comprised of these projection axes (10-20 dimensions is typically reported Churchland et al (2012); Cunningham and Yu (2014)). This low-dimensional state space is termed the neural state, which corresponds to coordinated action potential firing rates of the recorded neurons. This study reports results for various numbers of dimensions to show a more complete picture of the low-dimensional state and dynamics for unconstrained reaching and walking. Principal component analysis (PCA), an unsupervised mathematical model, and linear discriminant analysis (LDA), a supervised linear classification, were used to construct neural state spaces, and least-squares approximation modeled walking and reaching as linear dynamics systems.

### Principal component analysis

PCA was applied to multiple trials of neural data for each event type, and the latent spaces captured the highest-variance structure of neural activity. Before performing PCA, a Gaussian kernel with a 25 ms standard deviation was applied to the entire experimental session of 1kHz spike rasters for each of the 96 electrodes. The smoothed neural data was then normalized using the Standard Scaler method. All subsequent neural trials aligned to kinematic behavior had smoothed and normalized firing rates. To compare the dimensionality between swing and reach, PCA was performed separately on concatenated 200 ms trials for the swing and reach conditions, creating two different power spectra. The swing trials were randomly sampled to match the number of reach trials per day, displayed in Table S1. The dimensionality was computed by the number of principal components required to explain 80% percent of the variance, a common number reported in constrained experiments Churchland et al (2012); Yu et al (2009) (shown in Figure 2g and h). State-space trajectories of walking and reaching were constructed for each experimental day and the average of each trial was plotted for the top three dimensions (shown in Figure 2e and f).

### Neural subspace overlap analysis

Subspace overlap between gait and reach epochs were quantified using a similar method as in Elsayed et al (2016). PCA was performed separately for each of the five conditions (reach, swing, midswing, stance, midstance) for each day on 96-channel neural data of trials lasting 200 ms. The swing, midswing, stance, and midstance trials were randomly sampled to match the number of reach trials for a given day. In Figures 2e and f, the five conditions were projected onto the PCs of its own subspace, and the fraction of variance accounted for in the top 10, 20, and 30 PCs of a condition in its own subspace was calculated. The fraction of variance in a subspace was computed by taking the trace of the covariance matrix of the 96-channel spiking data after projecting it onto the top 10, 20, and 30 PCs. This was divided by the trace of the spiking covariance matrix to calculate the percentage of variance captured for each condition in its own subspace. The box and whisker plots in Figures 2e and f show the percent variance explained for the swing, midswing, stance, midstance, and reach conditions. To quantify the fraction of variance shared between the swing, midswing, stance, midstance, and reach epochs, the pairwise cross-projection of an epoch of one condition projected into the top 10 PCs of a different condition is shown in (Figure S6). Figure S6 shows the fraction of variance the walking and the reaching epochs contribute to two common subspaces: an 800 ms walking subspace, and a 1000 ms subspace with 800 ms of walking and 200 ms of reaching.

### Distance of gait trajectories from reach

The spatial distance in neural state space was quantified between the 200-ms reach trajectory and 200-ms segments of each gait phase (swing, midswing, stance, midstance). Figures 3e and f show the distribution of median distances in 10-dimensional space between each gait phase and reach trajectories for all trials. A rank sum test with bonferroni correction to adjust for multiple comparisons was performed on the means of the four histograms to determine if the distances were statistically significant between conditions. For both monkeys, p *<* 0.001 between swing vs. stance, swing vs. midstance, stance vs. midswing, and stance vs. swing, showing that the swing phases are significantly closer to the reach than the stance phases. Figure S7 show the same histograms, but for 40 and 96 PC dimensions. The results of the rank sum test for 40 and 96 dimensions were the same as in the 10-dimensional case. Figure S8a and b show the median distances between the reach and each gait phase from 0 to 200 ms for the top 10 PC dimensions. Figure S8e and f show the median distances from the reversed reach (200 to 0 ms). Figure S8c and d show the closest point of a gait phase tra-jectory for every point along the 200 ms reach trajectory for the top 10 PCs. Each individual line represents a different experimental day, and each point is color-coded by the closest gait phase for that millisecond.

### Fitting a linear dynamical system with least squares estimation

The low-dimensional neural trajectories for the gait cycle and reach were modeled using a least-squares approximation. Two separate subspaces were constructed using PCA each day consisting of 800 ms trials for the gait cycle and 800 ms trials for the reach for each day and were then modeled as a linear dynamics system using least-squares approximation. The least-squares approximation solves the linear matrix equation *Ax* = *y*, by minimizing the residual error ||*Ax* − *y*|| Boyd and Vandenberghe (2004). The problem statement for least-squares approximation is,

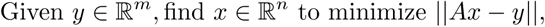

where, *A* is the dynamics update matrix, *x* is a vector of all the trajectories in neural state space for a given day, and *y* a vector of the differences between *x_t_*and time *x_t−_*_1_ for all the trajectories. The resulting solution gives the estimate of the next point of the neural trajectory *x_t_*_+1_ given all of the observations (*x*_0_*…x_t_*) from time 0 to t. Figures 3e and f visualize the walking (blue) and reaching (magenta) dynamics projected onto the walking dynamical system. The first real eigenvalue (squares) and the real component of the first complex eigenvalue pair (circles) of the dynamic A matrix for walking (*A_walk_*) and reach (*A_reach_*) quantified the dominant behavior of each system (Figures 3i and j). The real eigenvalue *λ* corresponds to an exponentially decaying or growing linear term, and the complex eigenvalue *λ* = *x* + *iy* corresponds to a decaying or growing sinusoidal term Boyd and Vandenberghe (2004). The fixed point of *A_walk_* was found by taking the Jacobian of the solution to the equation *ẋ*= *Ax* and is shown for two exemplar days per monkey in Figures 3e and f.

### Linear discriminant analysis (LDA)

LDA computes the largest hyperplane between the two scenarios by mapping the firing rates of neurons onto one latent space that represents the largest separation between two conditions while preserving the geometry of the neural activity. We performed LDA between the reach and swing conditions to find the projection that optimally separates these two conditions. The same number of 200 ms trials were concatenated for the swing and reach conditions for each experimental day. Trials were created by binning and summing every 25 ms for the 200 ms raster for each of the 96 electrodes. Swing trials were arbitrarily labeled as-1 and reach trials were labeled as 4. LDA was performed on all trials to create a linear decision boundary and decision function. In low-dimensional linear-discriminant (LD) space, the swing and reach are separable for both monkeys, shown in Figure S5a and b. Figure S5c and d show that reach and walk become more separable as the swing and reach evolve from 0 to 200 ms in LD1.

## Supplementary information

Table S1: Experimental sessions, number of laps, swing trial counts, and reach trial counts for Monkey U and Monkey C.

Table S2: Quantification of kinematics variables of the right wrist for the Monkey U.

Table S3: Quantification of kinematic variables of the left wrist for Monkey C.

Figure S4: Center of mass kinematics for three experimental days for Monkey U and Monkey C.

Figure S5: Neural activity for swing and reach are separable using LDA.

Figure S6: Percentage of variance explained relative to reach epochs, gait epochs, and two common subspaces.

Figure S7: Distance between reach and gait phase histograms for 40 and 96 dimensions.

Figure S8: Median distances from the reach phase to the four gait phases.

Figure S9: Monkey U heatmap of firing rates for all trials and electrodes.

Figure S10: Monkey U mean PSTH for across trial types and all electrodes.

Figure S11: Monkey C heatmap of firing rates for all trials and electrodes.

Figure S12: Monkey C mean PSTH for across trial types and all electrodes.

## Nonauthor Contributors

M. Wechsler, and M. Risch were responsible for animal care and surgical support. S.I. Ryu was responsible for non-human primate array implantation. I.E. Bray assisted in animal care.

## Acknowledgments

We thank K. Chin for administrative support.

## Author Contributions

ASL and MPS were responsible for collecting data with Monkey C (investigation). ASL, SEC, and EJJ were responsible for collecting data with Monkey U (investigation). MPS was responsible for developing the neural and camera data capture pipeline (data curation, methodology, software). ASL was responsible for developing the neural data filtering and synchronizing neural data (data curation, methodology, software). ASL was responsible for performing analysis (formal analysis) and data visualization. ASL wrote the original draft, the response to reviewers, and the updated draft. MUA developed the custom point-cloud hand-tagging pipeline in python, and ASL optimized it for this dataset (methodology). All authors contributed to hand-tagging point cloud frames (data collection). All authors reviewed and edited the manuscript. ASL and PN were responsible for data and analysis validation. PN was involved in all aspects of the project, conceptualized the work, supervised the effort, provided funding, and reviewed and edited the manuscript.

## Material and Correspondence

Please contact the corresponding author for correspondence and materials requests.

## Extended Data

**Table S1.** Experimental sessions, number of laps, and total swing trials, and total reach trial counts counts for Monkey U and Monkey C. A lap consists of 1-3 gait cycles and ended with a reach towards food in a bowl placed on the ground. Analysis for reach and all gait phases were trial matched to the number of good reaches for the contralateral arm for each experimental day. Monkey C had 9 experimental days and 633 analyzed out of 714 total reach trials from his right arm. Monkey U had 5 experimental days and 254 analyzed out of 272 total reach trials from his left arm.

| Monkey U | Laps | Swing trials | Reach trials | Monkey C | Laps | Swing trials | Reach trials |
| --- | --- | --- | --- | --- | --- | --- | --- |
| U221130_01 | 59 | 112 | 26 | C220405_01 | 62 | 172 | 16 |
| U221201_01 | 89 | 161 | 46 | C220406_01 | 50 | 128 | 20 |
| U221202_01 | 108 | 223 | 63 | C220407_01 | 62 | 96 | 24 |
| U221205_01 | 118 | 193 | 61 | C220411_01 | 148 | 451 | 82 |
| U221216_01 | 109 | 180 | 76 | C220414_01 | 225 | 476 | 160 |
|  |  |  |  | C220418_01 | 157 | 296 | 110 |
|  |  |  |  | C220419_01 | 202 | 380 | 156 |
|  |  |  |  | C220422_01 | 184 | 354 | 136 |

**Table S2.** Comparison of the swing and reach kinematics for the right wrist of Monkey U. The kinematic variables included the maximum velocity and peak time of the wrist, maximum acceleration, minimum acceleration, and start and end times of each trial. A Mood’s median test was performed on the median values of all the hand-tagged swing and reach trials to determine if there were any statistically significant differences. For Monkey U, no kinematic variables were statistically different with p *<* 0.05.

| Median values | Swing | Reach | Difference | p-value |
| --- | --- | --- | --- | --- |
| max velocity (m/s) | 0.23 | 0.22 | 0.012 | 0.18 |
| max acceleration (m/s<sup>2</sup>) | 0.0019 | 0.0018 | 0.00012 | 0.88 |
| min acceleration (m/s<sup>2</sup>) | -0.0019 | -0.0016 | -0.00020 | 0.15 |
| start time (ms) | 183 | 210 | -27 | 0.15 |
| peak time (ms) | 603 | 635 | -32 | 0.053 |
| end time (ms) | 827 | 847 | -20 | 0.39 |

**Table S3.** Comparison of the swing and reach kinematics for the left wrist of Monkey C. The kinematic variables are the same as in Table S2, and a Mood’s median test was performed for all hand-tagged trials comparing the swing and reach. Only the maximum velocity of the swing and reach showed a statistically significant difference with p *<* 0.05. All other variables showed no statistically significant difference.

| Median values | Swing | Reach | Difference | p-value |
| --- | --- | --- | --- | --- |
| max velocity (m/s) | 0.25 | 0.22 | 0.026 | 0.0028 |
| max acceleration (m/s<sup>2</sup>) | 0.0017 | 0.0017 | -0.000042 | 0.89 |
| min acceleration (m/s<sup>2</sup>) | -0.0014 | -0.0015 | 0.00011 | 0.20 |
| start time (ms) | 195 | 192 | 3 | 0.98 |
| peak time (ms) | 653 | 630 | 23 | 0.10 |
| end time (ms) | 810 | 818 | -8 | 0.48 |

---

**Fig. S4.**
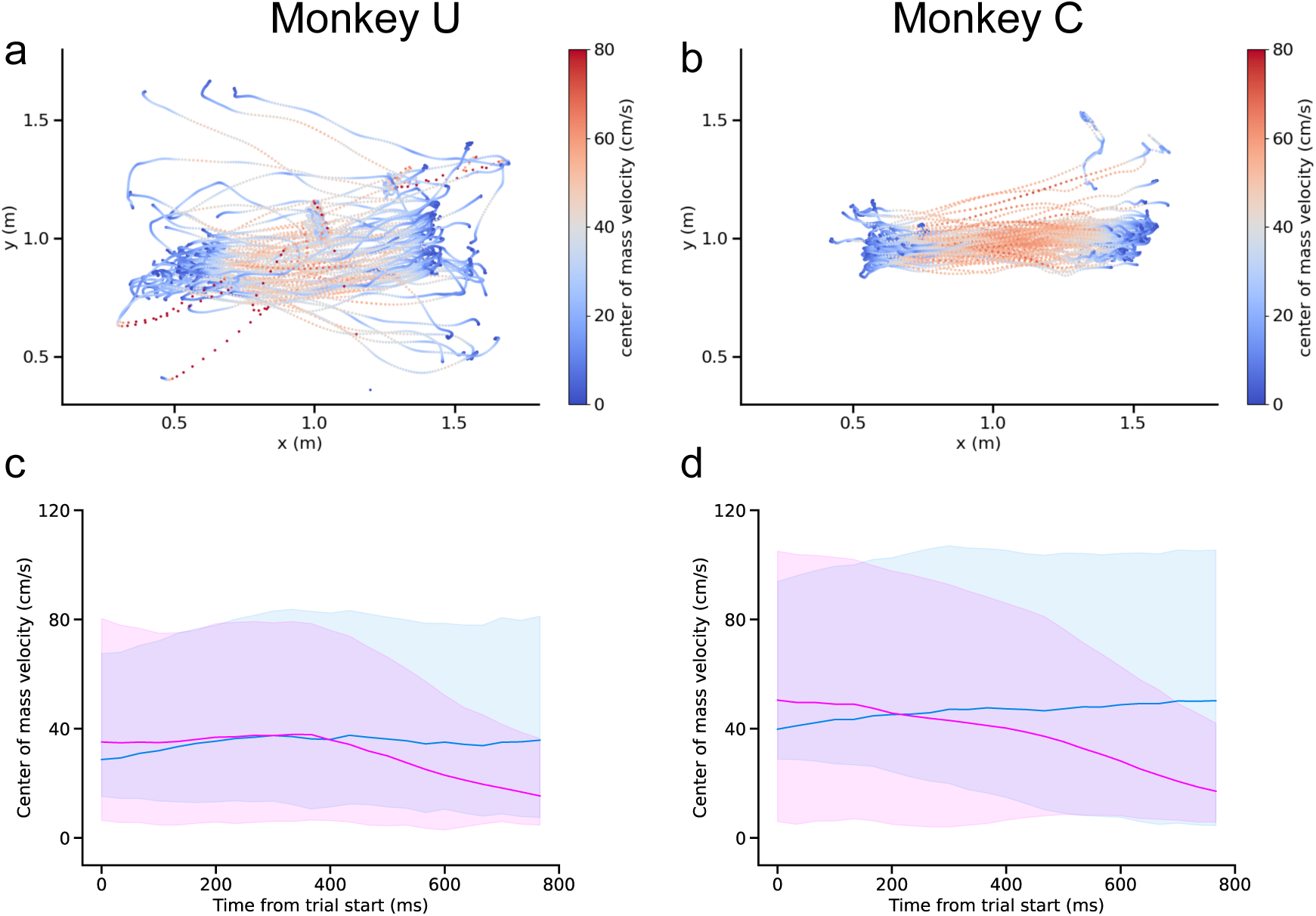
Center of mass trajectories in two dimensions as the monkey walked between two bowls for an example experimental day for Monkey U (a. U221201 01) and Monkey C (b. C220411 01). The kinematic center of mass was tracked by segmenting the monkey from the point cloud data and calculating the centroid of the resulting point cloud. Each dot represents the center of mass in meters of the monkey for a given point cloud frame, and the color scale represents the velocity at that time. Gait and reach trials were excluded if the center of mass position deviated from the typical walking trajectory between the two bowls. Monkey C reached a higher center of mass walking speed than Monkey U, as depicted by the higher intensity of red in the center of the kinematic trajectories. c) The average velocity of the center of mass during the 800 ms of swing (blue) and reach (magenta) is plotted for across all trials for Monkey U. Shading around the average line represents the 25% and 75% percentile. The trials times are aligned to the same trial times as in Figure 1e and f. The center of mass velocities were most similar between 200-400 ms representing the midstance epoch right before the beginning of the reach and swing phases. During the swing and reach phases (time 400-800), the center of mass velocity during the reach decreases while it stays constant during the swing. This is due to the monkey slowing down as he approaches the bowl for a goal-directed reach. d) Same as in c) but for Monkey C. The center of mass velocities during the reach and swing phases diverged than for Monkey U, possibly due to the higher center of mass walking velocities between bowls.

**Fig. S5.**
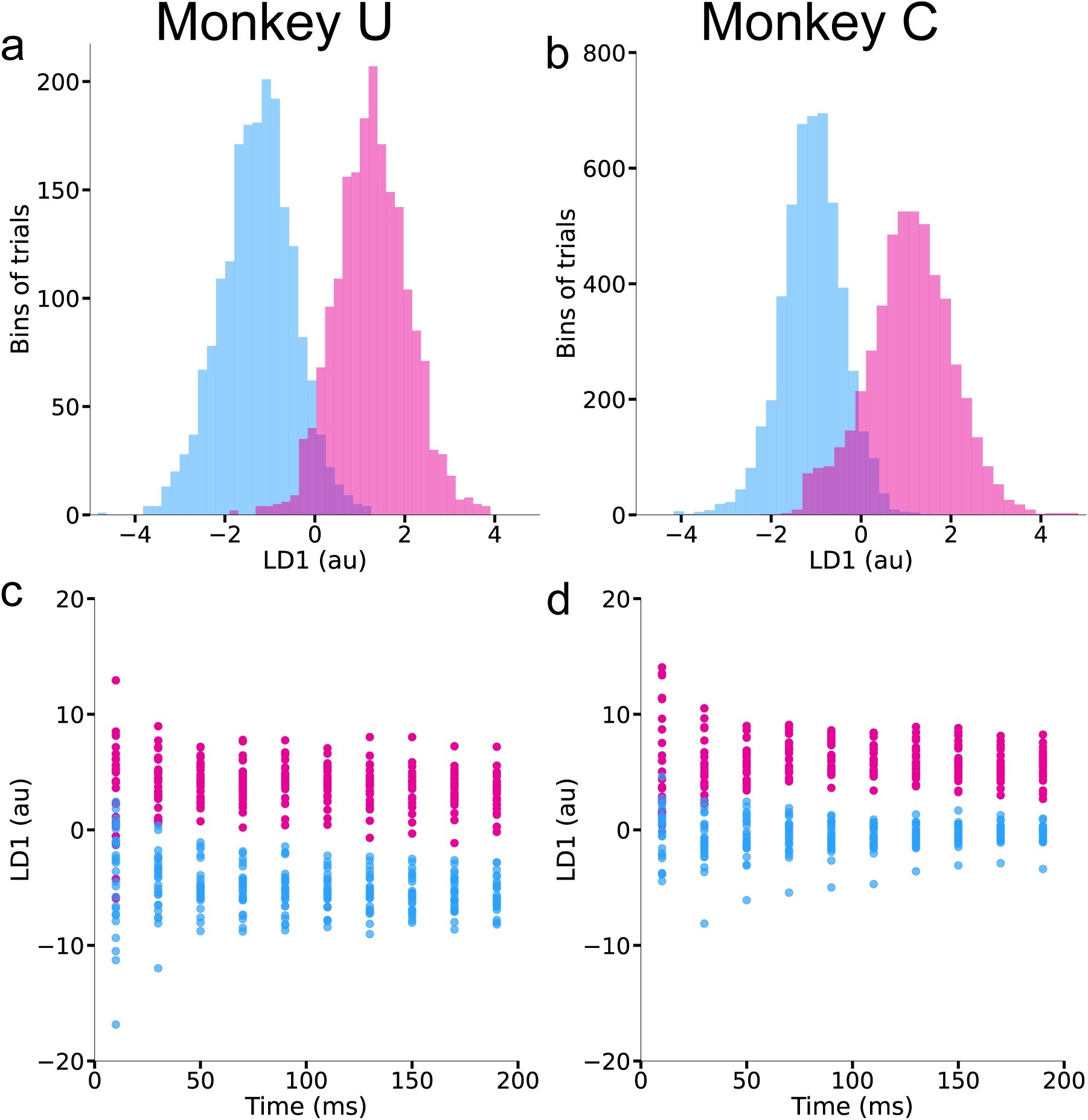
Neural activity for and unconstrained swing (blue) and unconstrained reach (magenta) are separable using LDA. Neural activity was concatenated for 200 ms trials binned every 25 ms for each swing and reach trial for a given day. a) LDA histogram of number of trials and bins for Monkey C showing a clear separation between the two conditions for all experimental days. b) LDA histogram for Monkey U. c) LDA trajectories for Monkey C. Trajectories were predicted using the linear decision boundary to classify unlabeled trials of 10 up to 200 ms bins every 20 ms. d) Same as in c) but for Monkey U.

**Fig. S6.**
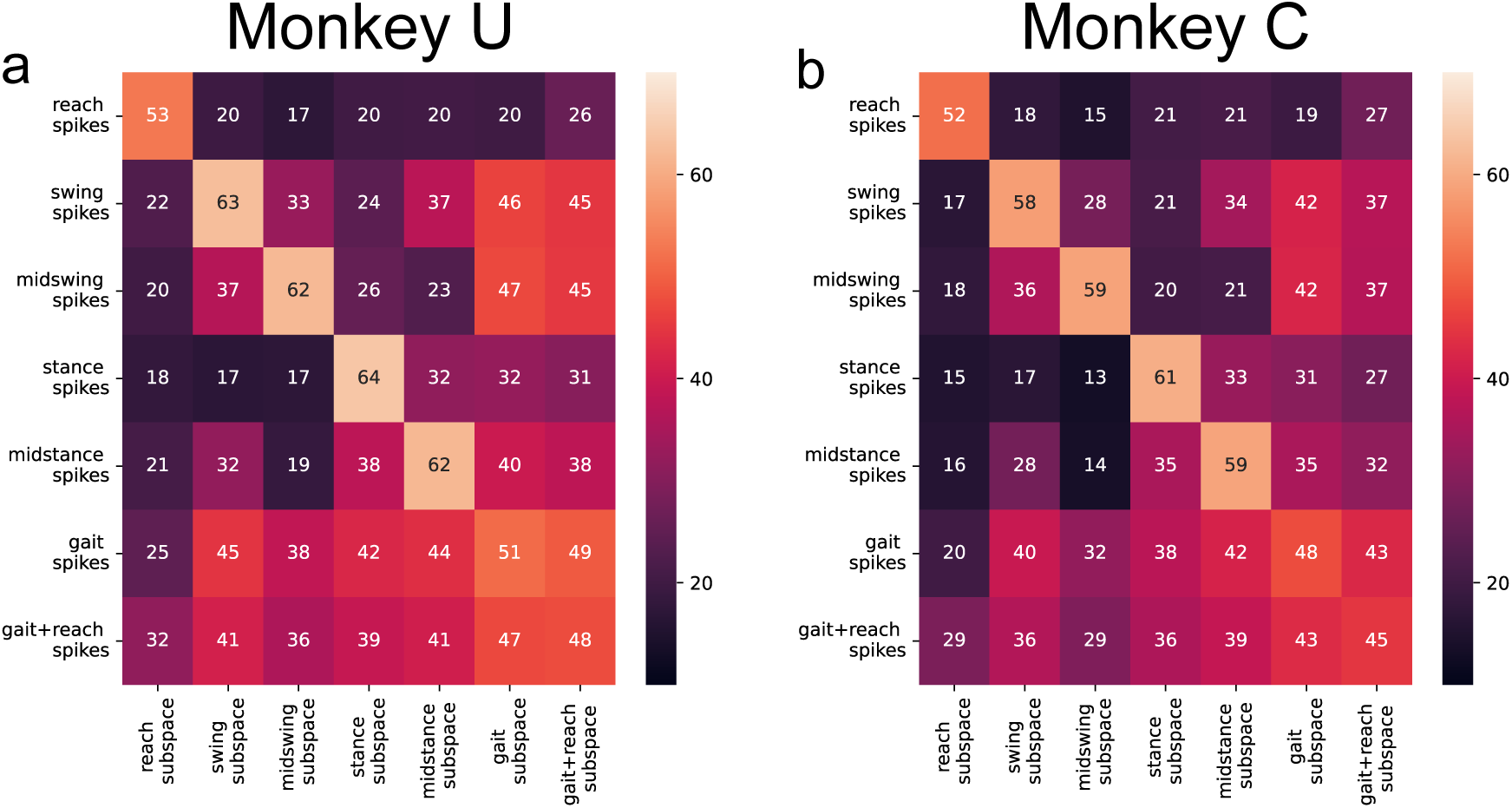
Percentage of variance explained for the reach, gait phases, and two common subspaces. Each entry in the matrix gives the fraction of variance accounted for as each epoch (reach, swing, midswing, stance, midstance, full gait cycle, gait plus reach) is projected into the top 10 PCs per epoch. The top ten reach PCs captured very little spiking gait data variance, and the top ten gait PCs captured very little reach spiking data variance. This finding reveals that reach occupies unique and non-overlapping subspaces relative to each gait epoch. Further, the top ten swing and midswing PCs captured very little spiking stance data variance, and the top ten stance PCs captured very little spiking swing and midswing data variance suggesting that swing and stance when separated may occupy different subspaces. When the swing and stance phases of the gait cycle were combined into a full gait PC subspace, the percentage of variance explained by reach spiking data was low, suggesting that reach has a low contribution to the state space of the full gait cycle. When each gait-epoch was projected onto the full gait cycle PC subspace, the amount of variance contributed by the two swing phases was higher than that of the two stance phases. However, the amount of variance for the four phases were comparable, between 30-40%. These results also hold true for one common subspace consisting of all four phases of the gait cycle and the reach. These results were consistent across two monkeys, Monkey U (a) and Monkey C (b).

**Fig. S7.**
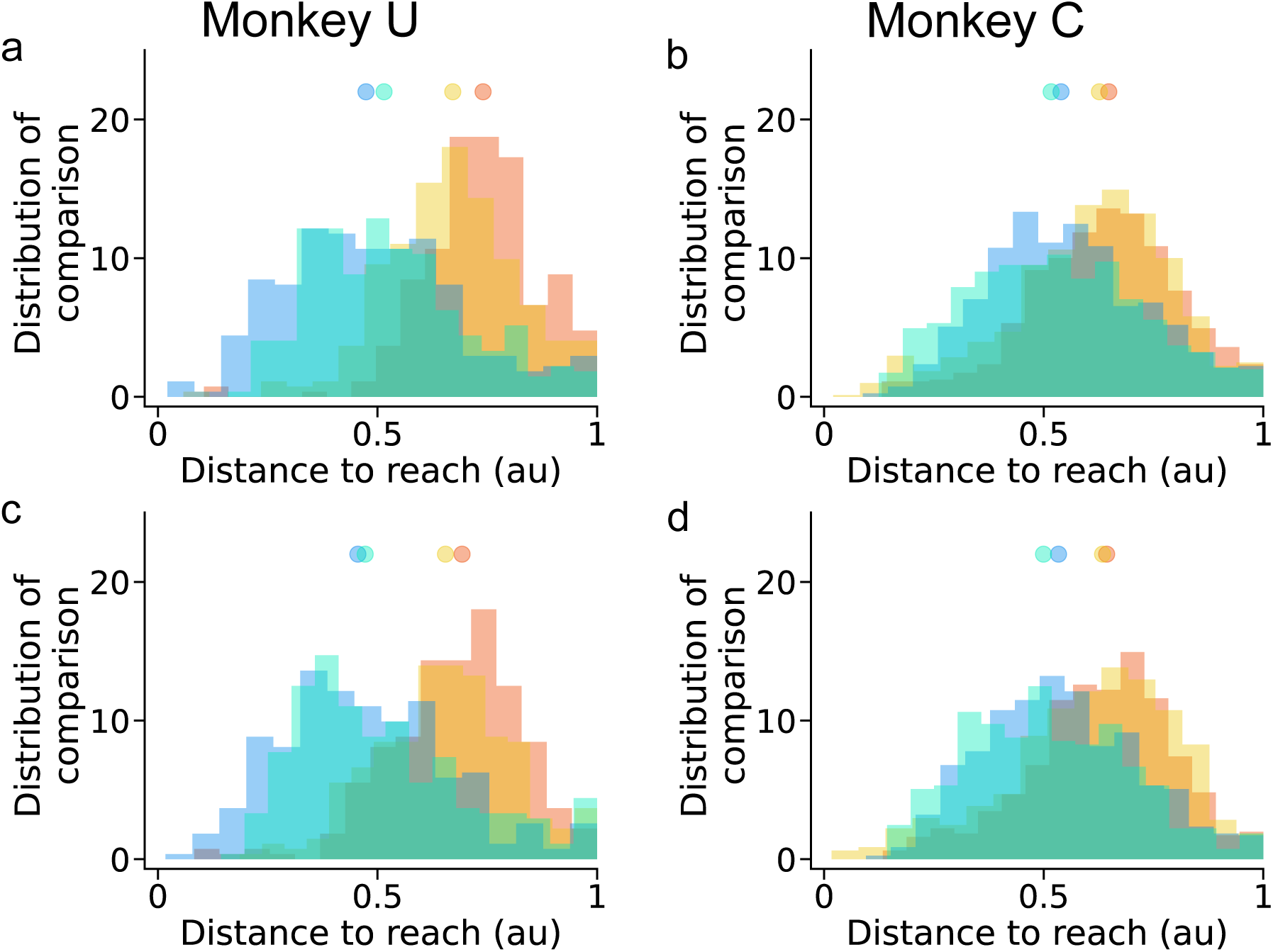
Same analysis as in Figure 2c and d for 40 and 96 dimensions. Similar distributions are observed as the 10-dimensional space after increasing the number of dimensions, as higher dimensions contribute proportionally smaller amounts of variance.

**Fig. S8.**
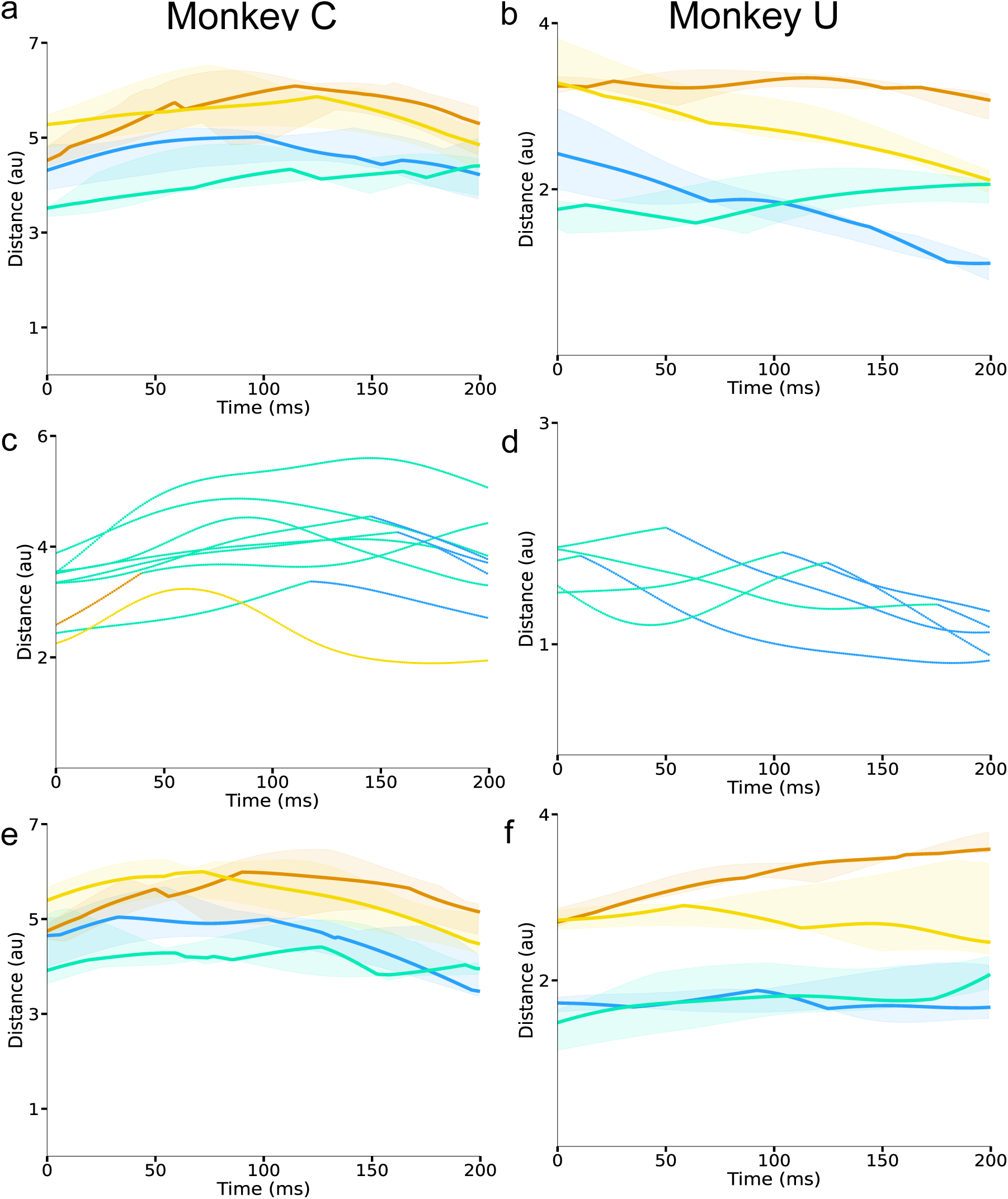
Median distances from the reach phase to the four gait phases, swing (blue), midswing (teal), stance (orange), and midstance (yellow). a) Median distance and interquartile ranges across experimental days from 0-200 ms of each gait phase to 0-200 ms of the reach. b) Same as in a) but for Monkey U. c) The closest median individual phase to each point of 0-200 ms median reach phase. Each line is a different experimental day. d) Same as in c) but for Monkey U. e) Median distance and interquartile ranges across experimental days from 0-200 ms of each gait phase to the reversed reach (200-0 ms). f) Same as in e) but for Monkey U.

**Fig. S9.**
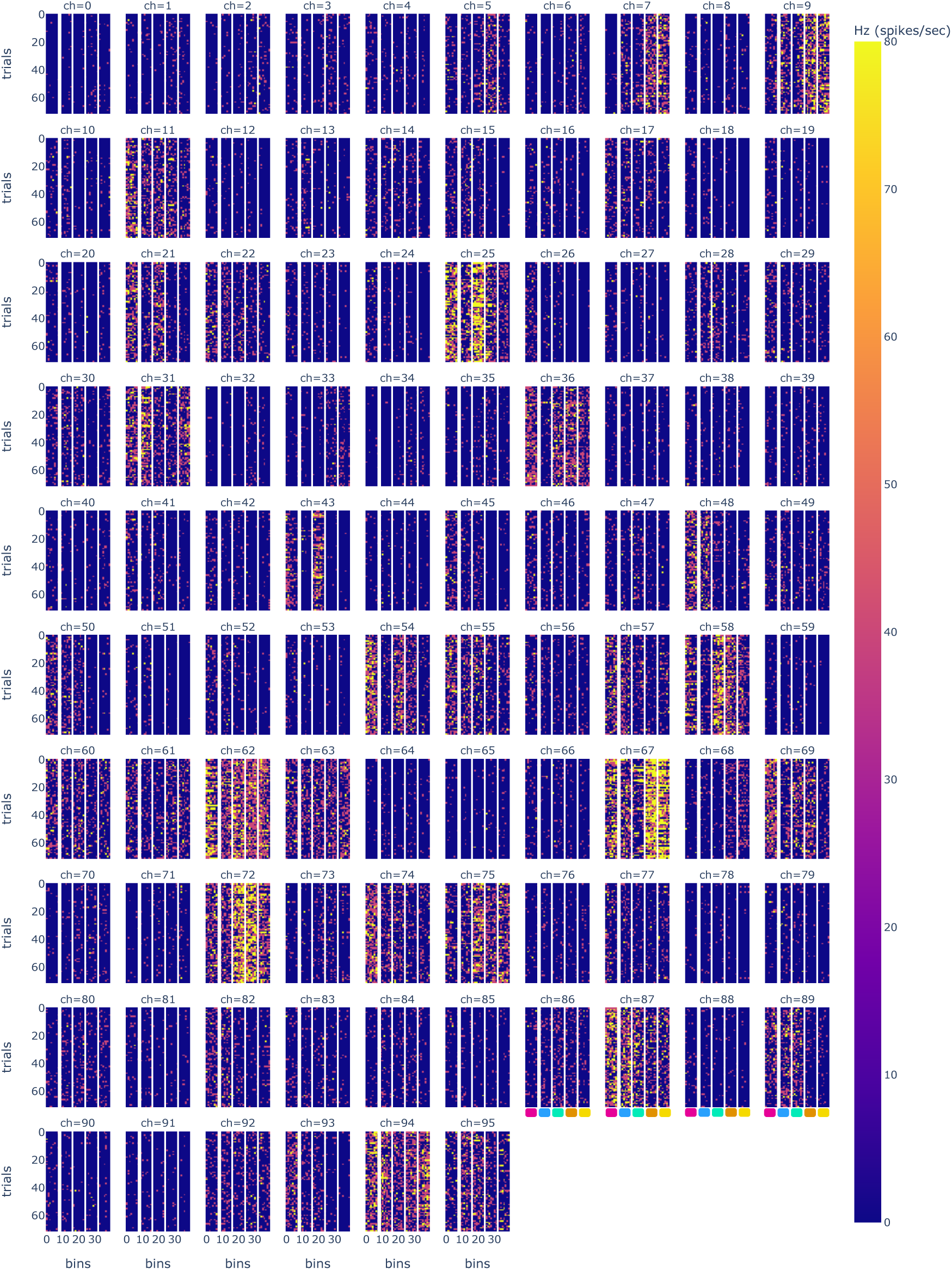
Heat map of firing rates for Monkey U, U221216_01 for all 96 electrodes. Panels are organized the same as in Figure S11

**Fig. S10.**
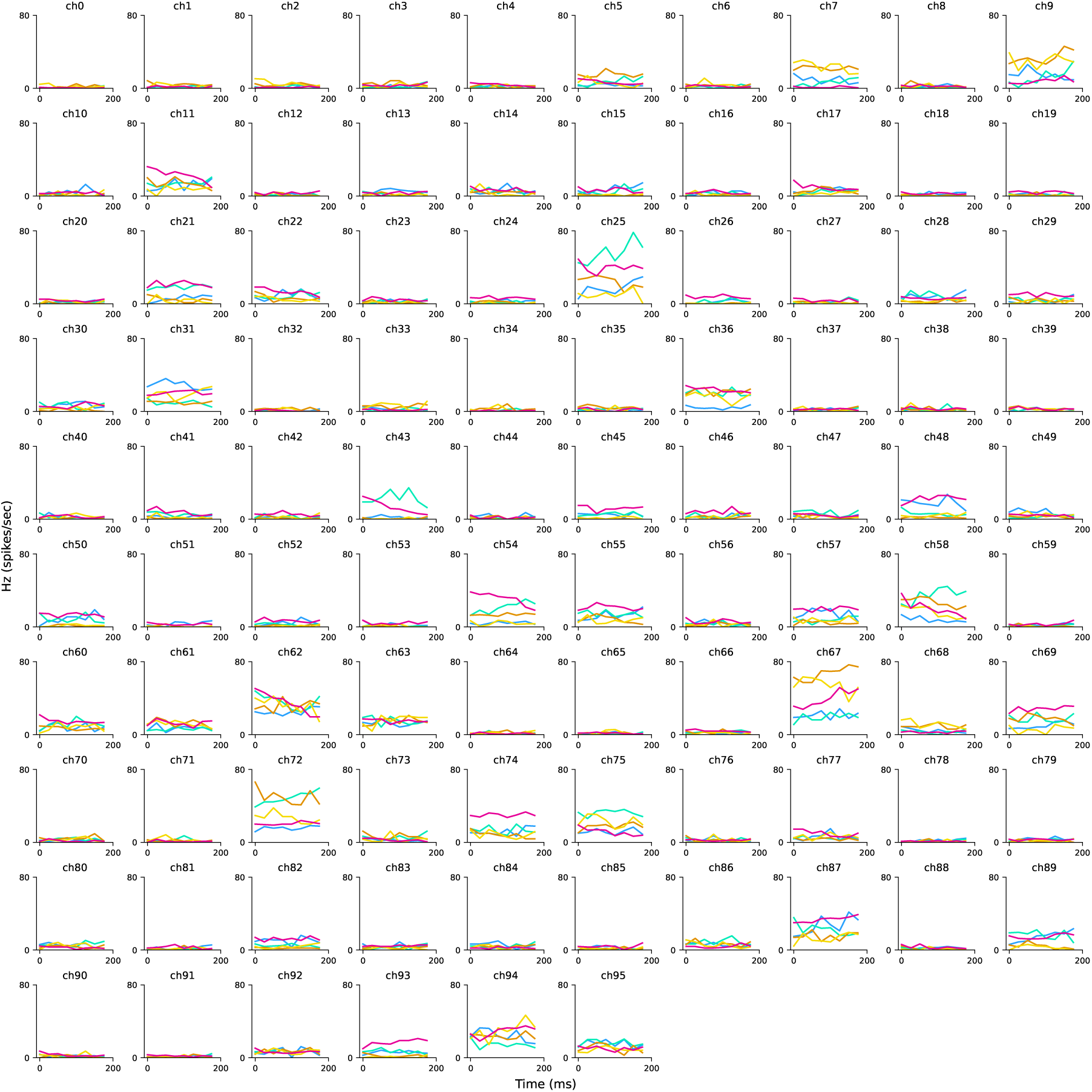
Mean PSTH for Monkey U for an example experimental day, U221216_01 for 96 electrodes from M1 array. Panels are organized the same as in Figure S12

**Fig. S11.**
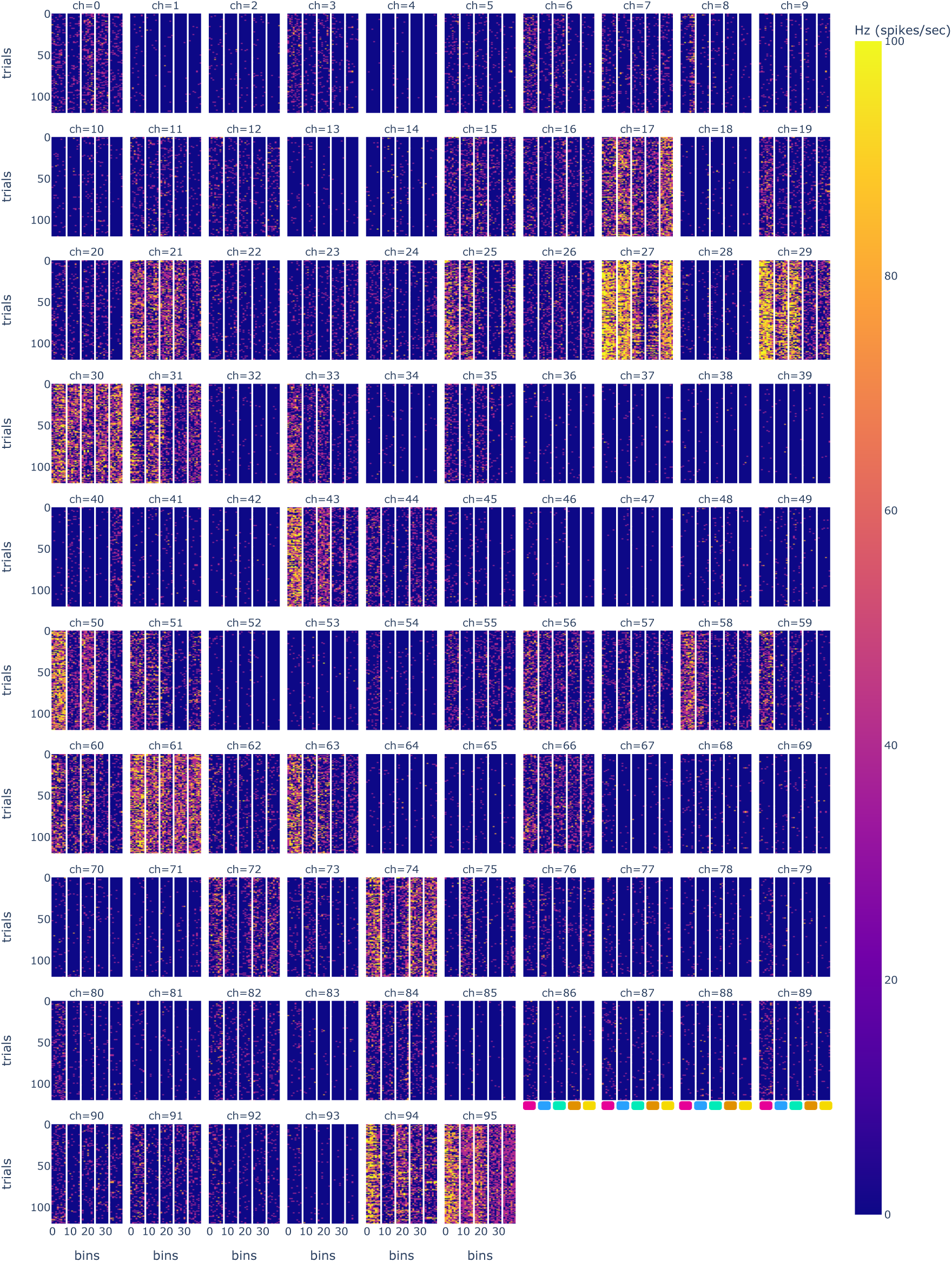
Heat map of firing rates for Monkey C, for an example experimental day C220422 01 for all 96 electrodes. Each panel within an electrode shows the firing rates for each trial across 8 bins of 25 ms for a total of 200 ms. Trials are ordered as reach, swing, midswing, stance, and midstance to directly compare reach and swing firing rates to each other.

**Fig. S12.**
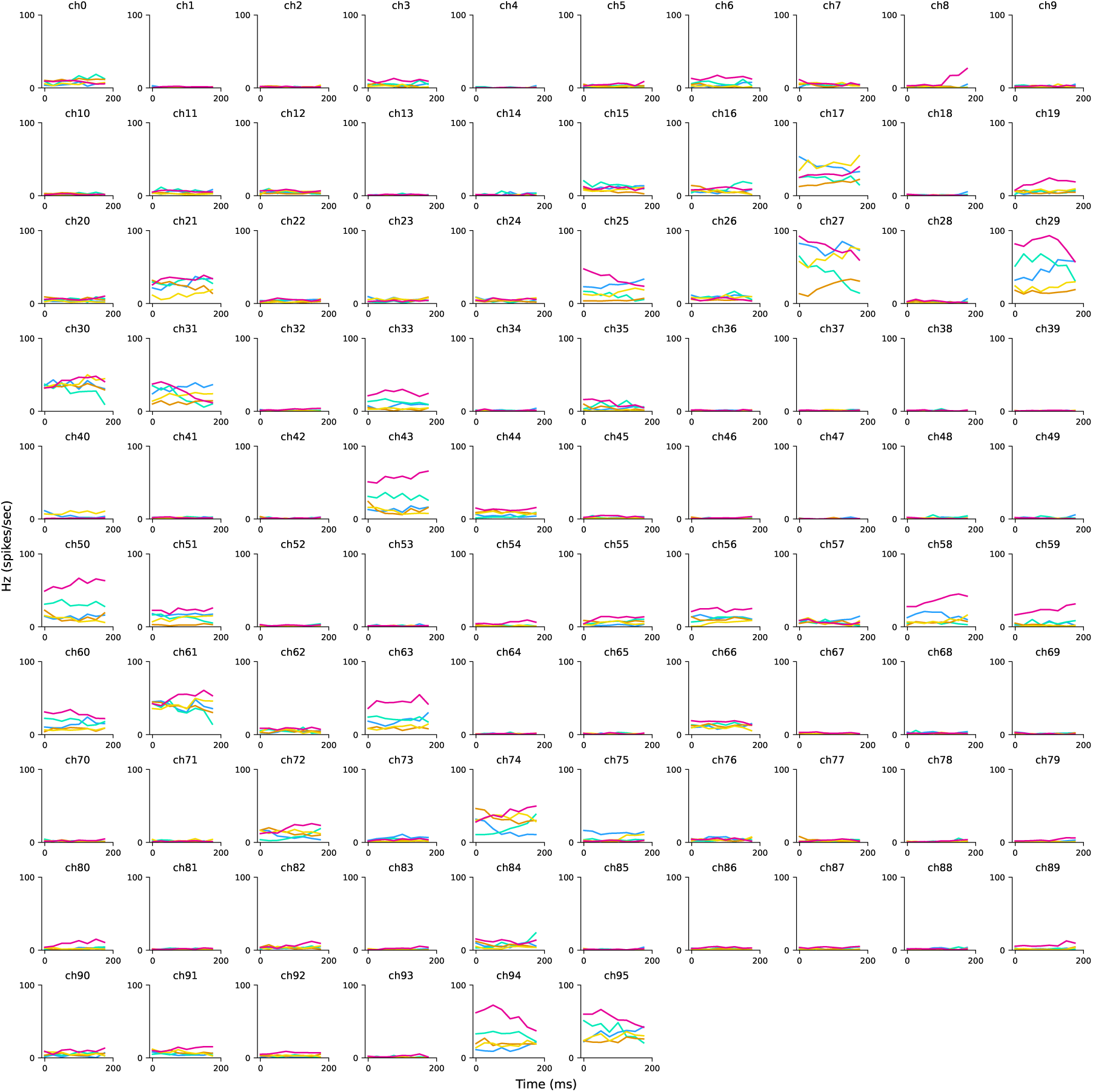
Mean peristimulus time histogram (PSTH) for Monkey C for an example experimental day C220422_01 for 96 electrodes from M1 array for the right arm. The mean firing rate across 200 ms was plotted for each trial, color coded the same as in Figure 1.

